# Nitrogen deposition is negatively related to species richness and abundance of threatened species in Swiss butterflies

**DOI:** 10.1101/2020.07.10.195354

**Authors:** Tobias Roth, Lukas Kohli, Beat Rihm, Reto Meier, Valentin Amrhein

**Affiliations:** Department of Environmental Sciences, Zoology, University of Basel, Basel, Switzerland; Hintermann Weber AG, Austrasse 2a, 4153 Reinach, Switzerland; Meteotest, Fabrikstrasse 14, 3012 Bern, Switzerland; Federal Office for the Environment (FOEN), Air Pollution Control and Chemicals Division, 3003 Bern, Switzerland

**Keywords:** Deposition model, elevational gradient, Lepidoptera, microclimate, microclimatic cooling, plant insect interactions, trophic interactions, vegetation

## Abstract

Nitrogen (N) deposition caused by agriculture and combustion of fossil fuels is a major threat to plant diversity, but the effects on higher trophic levels are less clear. In this study we investigated how N deposition may affect species richness and abundance (number of individuals per species) in butterflies. We started with reviewing the literature and found that vegetation parameters might be as important as climate and habitat variables in explaining variation in butterfly species richness. It thus seems likely that increased N deposition indirectly affects butterfly communities via its influence on plant communities. We then analysed data from the Swiss biodiversity monitoring program surveying species diversity of vascular plants and butterflies in 383 study sites of 1 km^2^ that are regularly distributed over Switzerland, covering a modelled N deposition gradient from 2 to 44 kg N ha ^−1^ yr^−1^. Using traditional linear models and structural equation models, we found that high N deposition was consistently linked to low butterfly diversity, suggesting a net loss of butterfly diversity through increased N deposition. At low elevations, N deposition may contribute to a reduction in butterfly species richness via microclimatic cooling due to increased plant biomass. At higher elevations, negative effects of N deposition on butterfly species richness may also be mediated by reduced plant species richness. In most butterfly species, abundance was negatively related to N deposition, but the strongest negative effects were found for species of conservation concern. We conclude that in addition to factors such as intensified agriculture, habitat fragmentation and climate change, N deposition is likely to play a key role in negatively affecting butterfly diversity and abundance.

**Article Impact Statement:** Nitrogen deposition negatively affects butterfly species richness and butterfly abundance, particularly in species of conservation concern.

**Data accessibility and reproducibility of results:** Data and R-scripts to reproduce the results of this manuscript including figures and tables are provided at https://github.com/TobiasRoth/NDep_butterflies. Raw data for analyses are provided in the folder “data”, and the folder “R” contains the R-Script that was used to export the data from the BDM database and to produce the figures and tables of the manuscript.

## Introduction

The increased flux of reactive nitrogen (N) in the biosphere and its deposition to ecosystems is considered as a major component of global change threatening biodiversity (Sala 2000). Increased N availability usually results in increased biomass production, in shifts in species composition (e.g., favoring eutrophic species), and often in a loss of plant species richness through competitive exclusion (Bobbink et al. 2010; Vellend et al. 2017). While the consequences of increased N availability are mainly documented for primary producers such as vascular plants (Bobbink et al. 2010), negative effects of increased N availability have also been found in species groups higher in the food chain, for example in insects (Haddad et al. 2001; WallisDeVries & van Swaay 2017).

Because plant and insect communities are closely linked, N-induced changes in plant communities are likely to induce changes in insect communities (Sassi et al. 2012). For example, because insects are often specialized on one or few plant species, the loss of plant diversity may negatively affect the diversity of insects (Knops et al. 1999; Haddad et al. 2001). Further, increased N availability favouring plant growth and biomass production is likely to alter the structure of the vegetation, thus leading to shifts in microclimatic conditions from open, dry and hot to more dense, humid and cool conditions, which will likely affect insects (WallisDeVries & vanSwaay 2006; WallisDeVries & van Swaay 2017).

However, our knowledge on how increased N availability affects consumer diversity is rather limited (Humbert et al. 2016; but see Haddad et al. 2001; WallisDeVries & van Swaay 2017). For example, in a literature review from 2016, only 18 (10%) of the 187 effect sizes on species richness reported from N-addition experiments were about invertebrates (Murphy & Romanuk 2016). Interestingly, the average effect size of those 18 studies suggests that the correlation between increased N availability and local-scale species richness of invertebrates is slightly positive (none of the 18 studies from this literature review investigated butterflies). Note, however, that since this review was published, the number of studies on N deposition effects on consumer diversity increased (WallisDeVries & van Swaay 2017; Schuldt et al. 2019).

Here, we complement the experimental studies on the effects of N deposition on higher trophic levels with an observational study using multiple field sites representing a large gradient of N deposition (i.e., a gradient study, Roth et al. 2017). The data are from the Biodiversity Monitoring Switzerland program (BDM) and contain information on species richness of vascular plants and butterflies in 383 study sites of 1 km^2^ and thus on the landscape scale, covering an N deposition gradient from 2 to 44 kg N ha^−1^ yr^−1^ (Roth et al. 2015). In previous studies, we found that high N deposition in these landscapes was associated with low values of six measures of plant diversity, including species richness (Roth et al. 2015). The BDM data thus provide an opportunity to examine possible direct and indirect effects of N deposition on species diversity of butterflies.

We started with a systematic literature review, searching for published studies investigating how butterfly species richness is related to environmental, land-use and vegetation parameters. The aim of the literature review was to compile a comprehensive list of predictor variables that could be important for explaining the variation in butterfly richness among our study sites. A second aim was to quantify how often N deposition was used as a predictor variable in such studies.

We then compiled the data from the BDM study sites and used traditional linear regression models to investigate how N deposition is correlated with butterfly species richness and how this correlation is affected depending on whether we accounted for all or only a selection of the other predictor variables. Since we assumed that a possible negative effect of N deposition on butterflies would be mediated by plant communities, we predicted that a negative effect would be weaker in models accounting for variables describing plant communities. We then used structural equation models (SEM) for examining the different paths by which environmental variables could affect butterfly species richness (Grace et al. 2010). In particular, we investigated how N deposition may negatively affect butterfly diversity via a negative effect on plant diversity (Topp & Loos 2018) and, additionally, via microclimatic cooling, for example because the increasingly productive and dense plant canopy may prevent caterpillars from absorbing solar radiation (WallisDeVries & vanSwaay 2006). Finally, we estimated the effect of N deposition on the abundance of the different butterfly species and examined how these effects differed between threatened species and species of less conservation concern.

## Methods

### Literature review

On 12 July 2019 we conducted a literature search in Web of Science. Our main aim was to compile a list of possibly relevant predictor variables for explaining butterfly species richness. We searched for original studies that applied some sort of multivariate regression models with several predictor variables and the variation in butterfly species richness among sites or grid cells as response variable. Since we aimed to quantify how often the different categories of predictor variables were used, we did not use specific search terms for nitrogen deposition or other predictor variables. Instead, we more generally searched for studies with titles that fulfilled the following search criteria: *[(butterfl* OR lepidoptera) AND (diversity OR richness)]; we* excluded studies with *[island OR tropic*]* in the title. Furthermore, the topic needed to contain *[“global change” OR driver* OR predictor OR variable].* See Appendix Al for a screenshot of the search setting in Web of Science. This search resulted in 95 studies. We excluded studies conducted in tropical rain forest and desert and found 32 studies that complied with our criteria (Appendix A2).

From the 32 studies, we extracted the predictor variables that were used to model butterfly species richness and assigned them to one of the following categories: (1) broad environmental category at the landscape level, including climatic gradients (from cool and humid to hot and dry) and climatic variability, and topographic variables (from low to high elevations; from northern to southern expositions; from low to high topographic variability); (2) habitat category at the level of habitat patches, including variables indicating the availability (from low to high total area of suitable habitat patches), configuration (from low to high suitability of habitat patch configuration) and diversity of habitat types (from low to high diversity of habitat patches) as well as land-use intensity (habitat patches with low to high land-use intensity); (3) vegetation category describing the vegetation or the conditions within the vegetation, including resource diversity (from low to high plant or flower richness) and micro-climate (from dense vegetation with cool and humid microclimate to open vegetation with hot and dry microclimate); and (4) other variables that did not fit (1) to (3), such as global vegetation index, area age or soil parameters.

For each of the first three categories (environmental, habitat and vegetation), we then grouped the predictor variables into sub-categories. The idea was that the predictor variables within a sub-category could be used to measure similar underlying processes that may affect butterfly diversity.

In Appendix A2 we present the entire list of predictor variables investigated in the 32 studies. For each study we extracted the investigated predictor variables and assigned the reported effect on butterfly species richness using the following coding: 1 = the effect of the category on butterfly diversity as measured by a predictor variable was positive; 0 = there was no obviously important effect; −1 = the effect of the category on butterfly diversity as measured by a predictor variable was negative; “interm” = the effect of the category on butterfly diversity peaked at intermediate level of the predictor variable. We coded an effect as important (1, −1 or “interm”) if the authors of the study mentioned in the abstract or discussion that they considered the reported effect size as important or relevant. If the authors did not make a statement about the importance of the reported effect, we judged the importance and direction of the effect ourselves, based on the reported point estimate and the precision (compatibility interval or standard error).

### Butterfly and plant data

We analysed the presence/absence of butterfly and plant species sampled between 2005 and 2009 in the Swiss Biodiversity Monitoring program (BDM, www.biodiversitymonitoring.ch). To monitor species diversity at the landscape scale, a sample grid of 428 equally-spaced study sites, each of 1 km^2^ size, was randomly selected. From the 428 study sites, seven sites with 100% water surface and 25 sites that were too dangerous for fieldwork because of their exposed alpine terrain were excluded a priori, resulting in 396 study sites.

Within each study site, surveyors walked along a 2.5 km transect that followed existing trails preferably near the diagonal of the grid cell (Plattner et al. 2004); the same transects were used to survey plants and butterflies. All transects were selected such that they lied entirely within the study sites. By using the existing trail network whenever possible, the location of the transects in the landscape was not random. As a consequence, the typical plant species of standing waters, marshes, and swamps were less fully represented than species of other major habitats (unpublished data).

The field protocol for butterflies was based on the British butterfly monitoring scheme (Pollard et al. 1995): transects were surveyed seven times between 21 April and 21 September in the lowlands, and four times between July and August above approximately 2000 m. The number of surveys corresponds to the shorter flying season of butterflies at higher elevations. The number of sites were selected such that sites at high and low elevations received approximately equal sampling effort per week of the flight season. During each survey, surveyors walked the transects in both directions and recorded all day-flying butterfly species (including *Hesperiidae* and *Zygaenidae)* within 5 m to each side of the transects on the way forth and back, respectively. Detectability varied by butterfly species and averaged 88% per survey (Kéry et al. 2009).

For the plant surveys, transects were surveyed by qualified botanists once in spring and once in summer, assuring that data collection spanned a large variation in flowering phenologies (Pearman & Weber 2007). At study sites with short vegetation period above approximately 2000 m, only one survey per field season was conducted. During each survey, surveyors recorded all plant species within 2.5 m to each side of the transects both on the way forth and back, respectively. The overall detection error was relatively small, with an average of 6.6% undetected presences per plant species as estimated in an earlier study using site-occupancy models (Chen et al. 2012).

Plant and butterfly surveys were usually conducted in the same years; each year, one fifth of the study sites were surveyed. Since we used the N deposition rates modelled for 2007 (see below), we selected the butterfly and plant data from the survey year that was closest to 2007 for each study site; this was the reason why survey data are from 2005 to 2009. In the analyses, we only included study sites for which both the plant and butterfly surveys met our standards of data collection or of weather conditions according to the protocol (Roth et al. 2014). This resulted in omission of 13 study sites and in a final dataset on plant and butterfly data collected in 383 study sites.

### Predictor variables

For all categories that were assigned to the predictor variables found in the literature review we included at least one predictor variable that was available for the BDM study sites (Table 1). Climate variables were extracted from the WorldClim database (Fick & Hijmans 2017). The source for the topographic data package was the GEOSTAT data base of the Federal Statistical Office (FSO, version 2006). Habitat data were derived from aerial data and are available on a grid with a 100-m resolution in the land-cover data package, also from the GEOSTAT data base of the FSO (version 2.0, 2013).

**Table 1:**
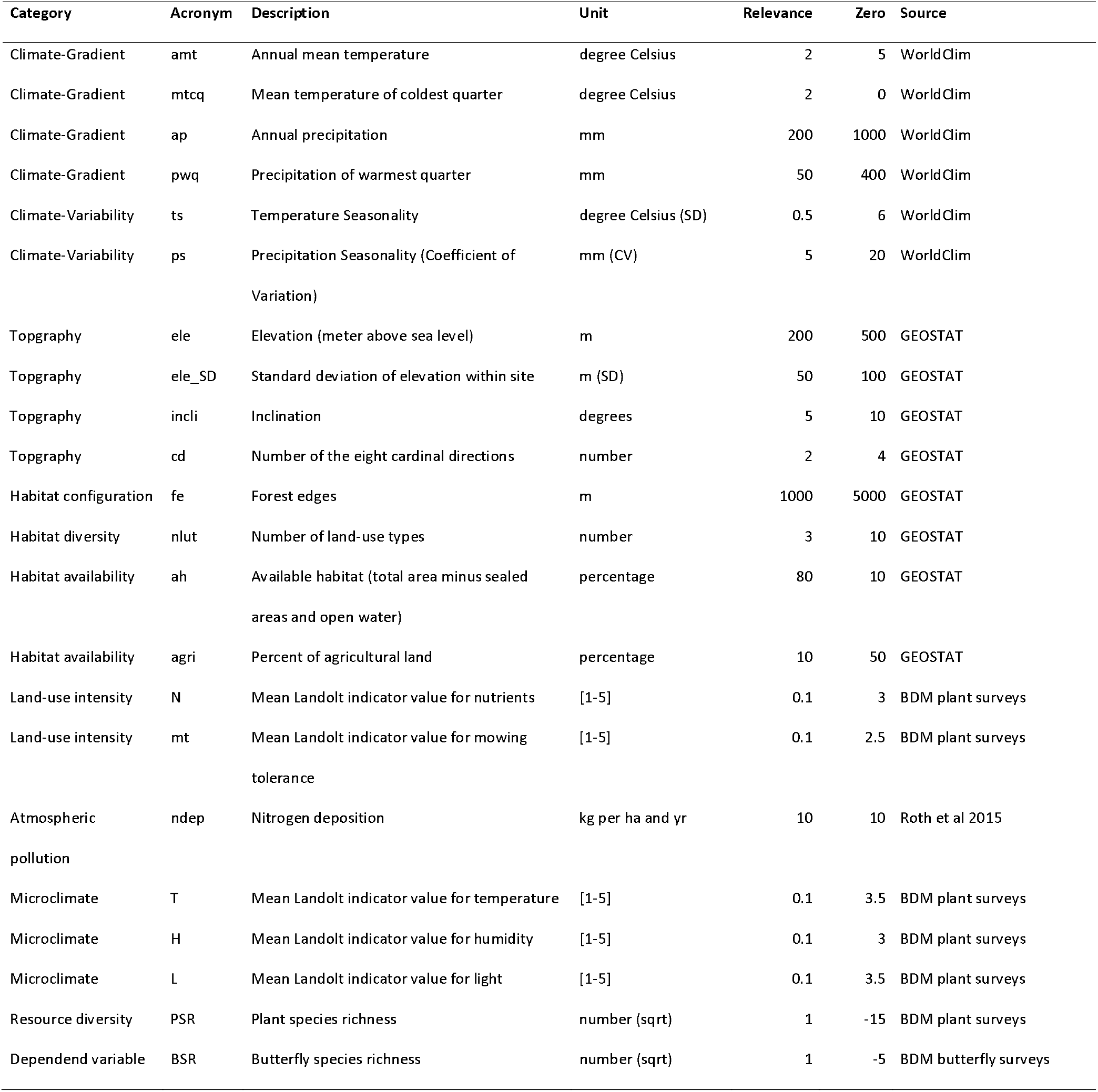
List of the predictor variables used to explain butterfly species richness. The categories are obtained from the literature review and reflect different processes that may affect butterfly diversity. For the statistical analyses we standardized the predictor variables by subtracting “Zero” and dividing it by “Relevance”. See main text for details.

Predictor values for land-use intensity and microclimate were derived from the species lists of recorded plants using Landolt indicator values that were developed for the specific situation in Switzerland (Landolt et al. 2010). Landolt values are ordinal numbers that express the realized ecological optima of plant species for different climate, soil or land-use variables. We used the mean Landolt indictor value of the recorded plant species for temperature and moisture as a measure for micro-climatic conditions in vegetation; we used Landolt indicators for nutrients and mowing tolerance as a measure for land-use intensity, and Landolt indicators for light as a measure of vegetation density (Table 1). Additionally, we used the total number of recorded plants as a measure of resource diversity.

N deposition was estimated for the year 2007 in 100×100 m grid cells across Switzerland, using a pragmatic approach that combined monitoring data, spatial interpolation methods, emission inventories, statistical dispersion models, and inferential deposition models (Roth et al. 2015; Rihm & Achermann 2016). For each study site of 1 km^2^, we averaged N deposition values from the cells containing parts of the transect used for the BDM surveys.

To obtain meaningful parameter estimates from the statistical models (see below), we centred the predictor variables by subtracting the value of the column “Zero” in Table 1 and standardized the predictor values by dividing them by the value of the column “Relevance”. Thus, the estimated intercept of the linear models is the predicted butterfly species richness for the values of the predictor variables as given in the column “Zero”. We chose these values to lie within the range of available data. The estimated slopes of the predictor variables indicate how much the butterfly species richness is changing if the predictor variable is increasing by the number that is given in the column “Relevance” in Table 1. To choose the number of the column “Relevance”, we asked ourselves the following question: what would be the minimum difference in the predictor value between two study plots that would result in detectable differences in, for example, vegetation. Although the choice of the value the in the column “Relevance” is arbitrary to a certain degree, it becomes easier to compare the parameter estimates among each other, which puts the focus on parameter estimates rather than on significance thresholds (Schielzeth 2010; Amrhein et al. 2019). A matrix with the scatterplots between all centred and standardized predictor variables is given in Appendix A3.

### Statistical methods

We used two different approaches for investigating the drivers of the spatial variation in butterfly species richness across Switzerland. The first approach is based on linear models, with the square root of butterfly species richness as response variable, and N deposition as focus variable included among the predictor variables in all tested models. Additionally, some of the other variables in Table 1 were included as covariates (i.e., additional predictor variables). We applied the following models: (1) full model that included the linear terms of all predictor variables in Table 1, (2) full model without microclimate variables, because microclimate is rarely considered in other studies on butterfly species richness, and climate and microclimate are usually correlated, (3) topo-climate model that included only the linear terms of the topography and climate-gradient variables, (4) climate model that included only the linear terms of the climate-gradient variables, (5) land-use model that included only the linear and quadratic term of elevation as a proxy for the climatic variation along the elevational gradient and the variables for habitat configuration, habitat diversity, habitat availability and land-use intensity, and (6) a minimalistic model that included only the linear and quadratic term of elevation as a proxy for climate and land-use intensity. All models assumed normal distribution of the residuals, and we examined this assumption for the full model using residual analyses. Model parameters were estimated in a Bayesian framework using the R-package arm (Gelman & Su 2018).

Our second approach was based on structural equation models (SEM, Hoyle 2012). We used the generic model that is given in Appendix A4 as a starting point for the analyses using SEM. In this generic model we assumed that butterfly species richness is mediated by vegetation structure and plant diversity (ovals and rectangles with grey background in Fig. A4). We further assumed that vegetation structure has an effect on plant diversity and, therefore, vegetation structure might affect butterfly diversity indirectly through its effect on plant diversity. Note that plant diversity is also likely to influence vegetation structure, and a bidirectional arrow between the two might have been more appropriate. Bidirectional arrows, however, are not possible to implement in SEMs.

Additionally, we assumed that different global change drivers such as climate, atmospheric nitrogen deposition or land-use intensity (white ovals) could each have independent effects on vegetation structure, plant diversity and butterfly diversity. While butterfly and plant species richness are measured variables in the BDM program (depicted in rectangles in Fig. A4), global change drivers and vegetation structure are latent variables that are measured by one or several of the predictor variables in Table 1. We present the results of different implementations of this generic model that vary in the number of global change drivers considered and in the selection of predictor variables used to measure the latent variables. Parameters of the SEMs were estimated using the R-Package lavaan (Rosseel 2012).

Finally, we tested for all butterfly species that were recorded in at least 20 study sites how the abundance of the species was related to N deposition. We used a generalized linear model with Poisson distribution with the number of recorded individuals of a species as dependent variable and the linear terms of all variables in Table 1 as predictor variables. We then compared the estimated effect size of N deposition between Red List species and the number of target species for which Swiss agriculture has particular responsibility of conservation (BAFU 2008; Wermeille et al. 2014).

## Results

### Literature review

From the 32 studies on butterfly species richness that we complied according to our inclusion criteria, we extracted the effect sizes of 252 predictor variables. Predictor environmental variables were included in 75% of studies, habitat variables were included in 84% of studies and vegetation variables were included in 47% of studies. Nitrogen deposition was considered in none of the compiled studies.

While predictor variables for the vegetation category were less likely to be considered in these studies compared to environmental and habitat variables, their importance (as estimated by the proportion of times the variables were considered important) was similar to the importance of the predictor variables for the environmental and the habitat category (Table 2). Furthermore, resource diversity of the vegetation was the variable with the most consistent effect (regarding the direction of the effects) across all variables considered in the reviewed studies (Table 2).

**Table 2:** Summary of the literature review. Given are for each subcategory the number of different indicator variables, the number of studies, the proportion of times an “important” effect was identified, and the direction of the important effect size (the mean direction of effects not coded as “zero”, excluding effects that peaked at intermediate levels).

| Category | Subcategory | Number of variables | Studies | Importance | Direction |
| --- | --- | --- | --- | --- | --- |
| Environment | Climate-Gradient | 10 | 18 | 0.63 | 0.26 |
| Environment | Climate-Variability | 4 | 8 | 0.56 | 0.00 |
| Environment | Topography | 3 | 13 | 0.45 | 0.19 |
| Habitat | Habitat configuration | 6 | 7 | 0.58 | -0.10 |
| Habitat | Habitat diversity | 4 | 11 | 0.64 | 0.27 |
| Habitat | Habitat-availability | 11 | 20 | 0.64 | 0.15 |
| Habitat | Land-use intensity | 25 | 19 | 0.57 | -0.42 |
| Vegetation | Microclimate | 3 | 7 | 0.64 | 0.11 |
| Vegetation | Resource diversity | 8 | 13 | 0.59 | 0.57 |
| Others | — | 4 | 6 | 0.29 | 0.00 |

### Field study

Based on the linear models that we applied to the BDM data, we found that butterfly diversity decreased with increasing N deposition. The amount of this negative effect of N deposition on butterfly species richness was similar for all considered models (Fig. 1). Except for the climatic variables (annual mean temperature, mean temperature of coldest quarter of the year, and temperature seasonality), N deposition was the variable with the highest absolute effect size in the full model (Table 3).

**Figure 1:**
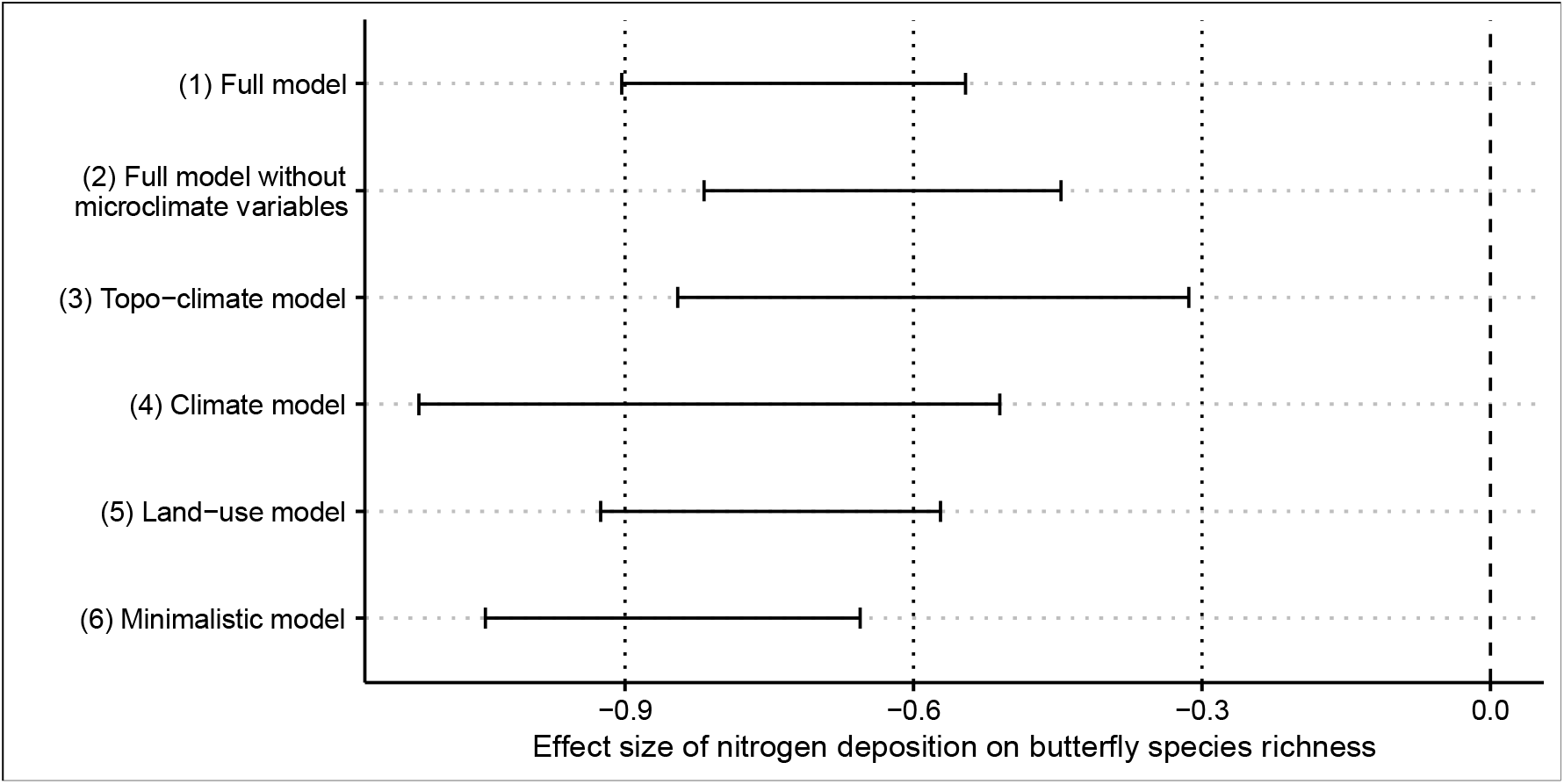
Effect sizes (with 95%-compatibility intervals; Amrhein et al. 2019) of nitrogen deposition on the square root of butterfly species richness based on linear models that differ in selection of predictor variables. See statistical methods for a description of the different models.

**Table 3:** Parameter estimates of the full model that explains the butterfly species richness with the linear terms of all the variables listed in Table 1. Ordering of the variables according to the absolute value of the estimate.

| Predictor variable | Description | Estimate | Std-Error | P-value |
| --- | --- | --- | --- | --- |
| mtcq | Mean temperature of coldest quarter | 2.405 | 1.032 | 0.020 |
| amt | Annual mean temperature | -2.179 | 1.072 | 0.043 |
| ts | Temperature Seasonality | 1.341 | 0.475 | 0.005 |
| ndep | Nitrogen deposition | -0.719 | 0.094 | <0.001 |
| ele | Elevation (meter above sea level) | 0.429 | 0.090 | 0.000 |
| PSR | Plant species richness | 0.248 | 0.031 | <0.001 |
| pwq | Precipitation of warmest quarter | 0.236 | 0.105 | 0.026 |
| T | Mean Landolt indicator value for temperature | 0.200 | 0.045 | <0.001 |
| N | Mean Landolt indicator value for nutrients | -0.198 | 0.060 | 0.001 |
| ele_SD | Standard deviation of elevation within site | -0.193 | 0.102 | 0.059 |
| L | Mean Landolt indicator value for light | -0.167 | 0.031 | <0.001 |
| incli | Inclination | 0.165 | 0.064 | 0.010 |
| ap | Annual precipitation | -0.113 | 0.124 | 0.364 |
| agri | Percent of agricultural land | 0.091 | 0.021 | <0.001 |
| ps | Precipitation Seasonality (Coefficient of Variation) | -0.081 | 0.061 | 0.188 |
| ah | Available habitat (total area minus sealed areas and open water) | 0.058 | 0.033 | 0.082 |
| fe | Forest edges | 0.030 | 0.012 | 0.018 |
| nlut | Number of land-use types | 0.028 | 0.051 | 0.580 |
| mt | Mean Landolt indicator value for mowing tolerance | 0.018 | 0.054 | 0.738 |
| cd | Number of the eight cardinal directions | -0.017 | 0.043 | 0.690 |
| H | Mean Landolt indicator value for humidity | -0.016 | 0.043 | 0.712 |

The results of the structural equation model (SEM) that included climate, N deposition, land-use intensity and habitat availability as global change gradients potentially affecting vegetation structure, as well as plant species richness and butterfly species richness are given in Fig. 2a. Based on the results of this model, butterfly species richness was affected (in descending order of the absolute value of the effect sizes) by climate (highest butterfly richness in warm and dry climate; estimate ± SE: 0.50 ± 0.054), plant species richness (butterfly richness increasing with plant richness; 0.38 ± 0.025), and microclimate (higher butterfly richness in areas with warm, dry and open vegetation than in areas with closed and humid vegetation; 0.13 ± 0.041).

**Figure 2:**
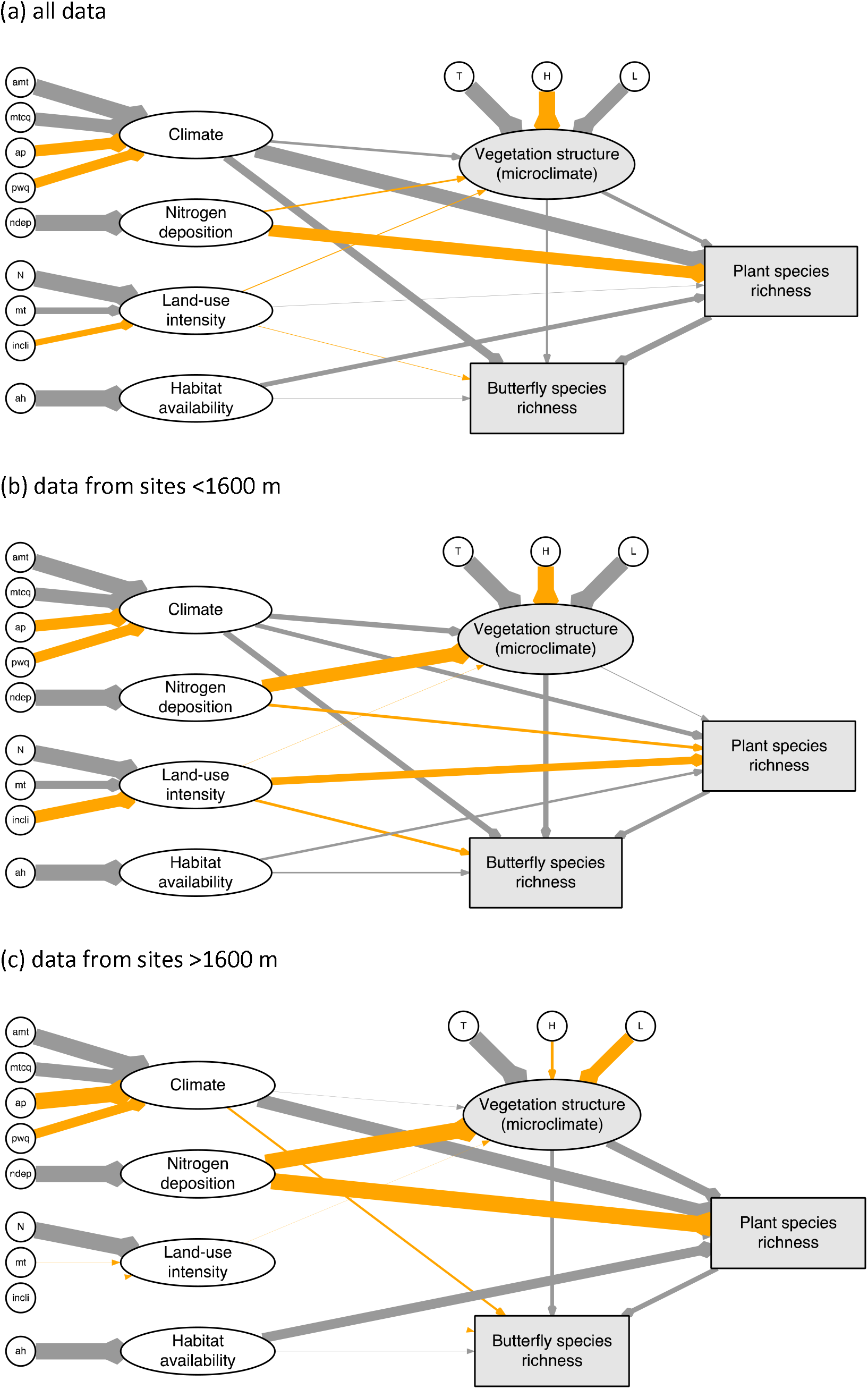
Results obtained from applying the structural equation model to (a) all data from the Swiss Biodiversity Monitoring program, (b) the data of the sites below 1600 m, and (c) the data of the sites above 1600 m. Depicted are presumed effects of different global change drivers and vegetation variables on butterfly species richness at 1-km^2^ resolution in Switzerland; the thickness of the arrows is relative to the absolute values of the effect sizes, with arrows in grey indicating positive effects, and arrows in orange indicating negative effects.

The effects of land-use intensity and habitat availability on butterfly species richness seemed to be rather weak (land-use intensity: −0.035 ± 0.021; habitat availability: 0.026 ± 0.025). However, at sites below 1600 m, where land-use is usually intense, the negative effect of land-use intensity on butterfly diversity was much stronger (−0.19 ± 0.034, Fig. 2b).

The results of applying the SEM to the data of all sites further suggest that higher N deposition rates lead to denser, more humid and cooler microclimates within the vegetation (−0.10 ± 0.042) and to lower plant species richness (−0.73 ± 0.15, Fig. 2a). At elevations below 1600 m, N deposition mainly affected vegetation structure (vegetation structure: −0.85 ± 0.097; plant species richness: −0.14 ± 0.17, Fig. 2b), and the negative effect of N deposition at higher elevations was strong both regarding vegetation structure and plant species richness (vegetation structure: −2.43 ± 1.011; plant species richness: −2.11 ± 1.09, Fig. 2c). Note, however, that the latent variable vegetation structure has a different meaning below and above 1600 m, as the mean Landolt indicator value for light (L) is positively correlated below 1600 m (Fig. 2b) and is negatively correlated above 1600 m (Fig. 2c). This makes intuitive sense, as below 1600 m the coldest habitats are the shaded ones within the forest, while above 1600 m the coldest habitats are the open habitats with hardly any vegetation that would protect from freezing temperatures.

From the 183 butterfly species that were recorded, 113 (62%) species were recorded in at least 20 study sites. The abundance (number of recorded individuals) of most of these 113 species decreased with increasing N deposition as revealed by generalized linear models that were applied to each species separately. The negative effect was strongest for near-threatened and vulnerable species (near-threatened: 24 species; vulnerable: 3 species; note that no critically endangered or endangered species were among the 113 analysed species), intermediate for the target species for which Swiss agriculture has particular responsibility of conservation (58 species including 19 of the near threatened and 2 of the vulnerable species) and weakest for the remaining species (Fig. 3). The estimated effect size for N deposition for each species is given in Appendix A5.

**Figure 3:**
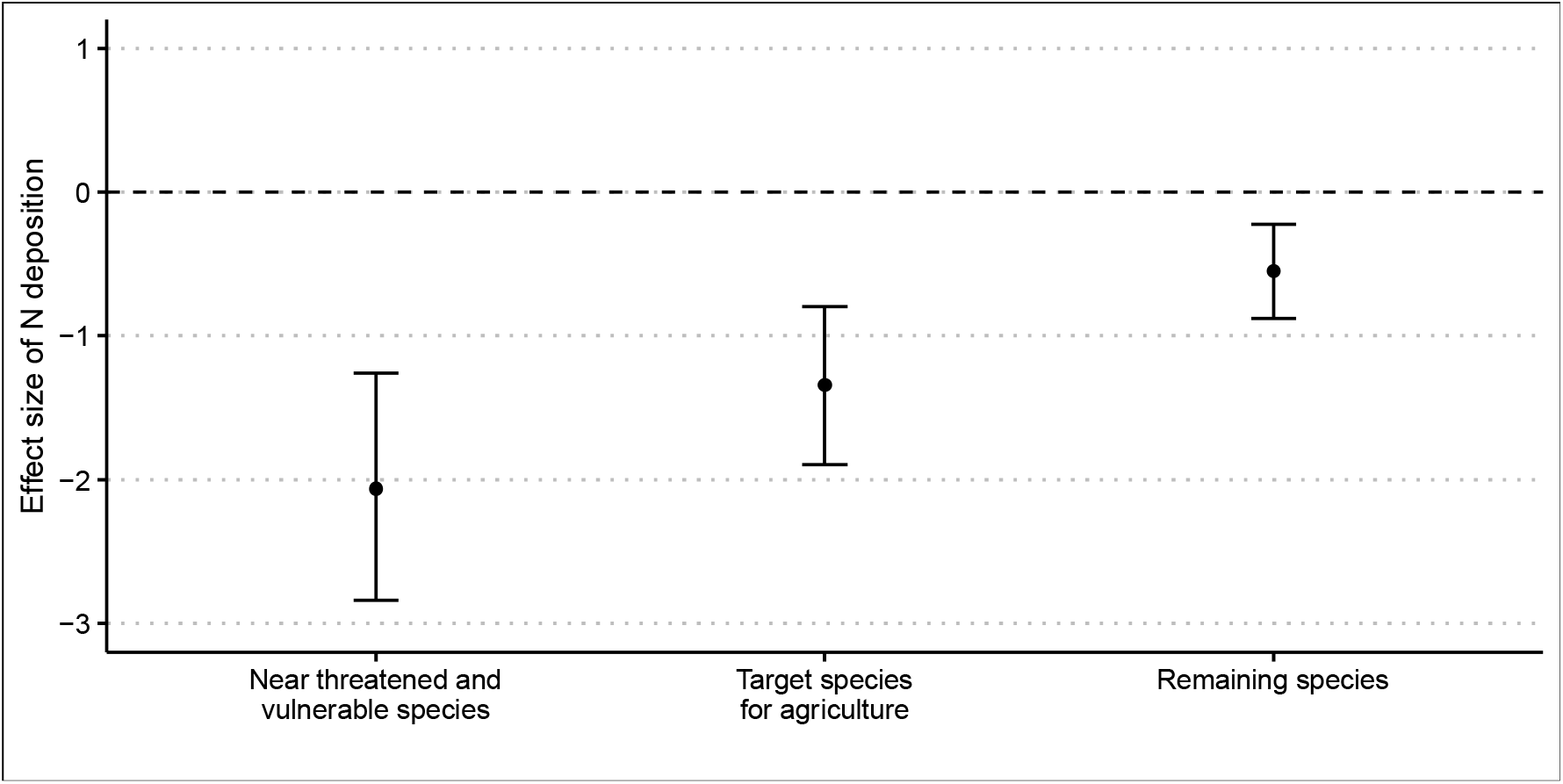
Average effect sizes of N deposition on butterfly abundance (number of recorded individuals) for all species with at least 20 records. The averages are given separately for near-threatened or vulnerable species, for target species for which Swiss agriculture has particular responsibility of conservation, and for the remaining species. The lines are 95%-compatibility intervals.

## Discussion

### Nitrogen deposition effect on butterflies and its conseguencesfor conservation

Our results confirm the importance of Nitrogen (N) deposition as a largely negative driver of butterfly species richness and abundance (number of recorded individuals) in Switzerland. Previous studies have found that N deposition affects butterfly species differently depending on their preferred food plant or other factors (WallisDeVries & vanSwaay 2006). For example, population sizes of butterfly species that depend on nutrient-poor conditions tend to decrease with increasing N deposition, while population sizes of species that depend on nutrient-rich conditions, or N-favoured plant species, tend to increase (Öckinger et al. 2006; Betzholtz et al. 2013). Our study complements these results by suggesting that species of conservation concern are particularly affected by N deposition, and at the landscape scale, species-dependent N-deposition effects sum up to a net loss of butterfly species richness due to increased N deposition.

None of the studies that we compiled in our literature review included N deposition as a predictor variable for butterfly species richness. Although our literature research was not exhaustive, the absence of N deposition in the 32 reviewed studies suggests that the negative effect of N deposition on butterfly communities has probably been underestimated so far. Given the global insect decline in terrestrial ecosystems (van Klink et al. 2020), the lack of awareness of N deposition as a negative driver of insect populations seems particularly relevant, because promising strategies to mitigate or even reverse the negative trends in insect populations might be overlooked. For example, mowing with biomass removal or grazing, which both remove large amounts of N, may have a positive effects on butterfly diversity, at least if the intensity of mowing or the density of grazers is not too high (Jones et al. 2017). Indeed, in a recent review on the factors believed to be responsible for the observed collapses of insect populations, Wagner (2020) concludes that “the potential consequences of atmospheric nitrogen deposition are grave and worthy of greater attention”.

### Mechanistic links between Nitrogen deposition and butterfly communities

Given that negative effects of increased N deposition on plant communities are well established (e.g. reduced plant diversity or increased vegetation density; Bobbink et al. 2010; Vellend et al. 2017), negative effects of increased N deposition on butterfly diversity through its effects on plant communities are likely to occur (Schuldt et al. 2019). At least three main mechanisms have been proposed: First, reduced plant diversity due to increased N deposition could result in reduced food diversity for butterflies (Zhu et al. 2016). Second, increased N deposition resulting in higher plant biomass and denser vegetation could lead to microclimatic cooling that, for example, may prevent caterpillars from absorbing solar radiation to attain optimal body temperatures (WallisDeVries & vanSwaay 2006). Third, the chemical composition of plants could change due to increased N deposition, resulting in reduced food plant quality (Habel et al. 2016). Note, however, that for some species a higher nitrogen level of host plants may also increase food quality for larvae (Pullin 1987).

The results of the structural equation models applied to the Swiss Biodiversity Monitoring (BDM) data seem to support the first two pathways: Particularly at higher elevations, where the negative effect of land-use intensity on plant species richness was reduced, increased N deposition was correlated with reduced plant species richness, and plant species richness was positively related to butterfly species richness (Fig. 2c). This suggests that at higher elevations, the negative effect of N deposition on butterfly species richness is mediated by its negative effect on plant species richness. At lower elevations, N deposition was mainly correlated with denser vegetation (i.e., with plant indicator values that are associated with less light) and with cooler and more humid vegetation, which was correlated with lower butterfly species richness. This suggests that at lower elevations, microclimatic cooling through increased N deposition contributes to a reduction in butterfly species richness (Fig. 2b).

We do not have data to directly investigate the third explanation stating that decreased butterfly diversity due to N deposition could be caused by reduced food plant quality. However, when we allowed for a direct effect of N deposition on butterfly species richness in the structural equation model (Appendix A6), the results suggested a quite strong negative effect of N deposition on butterfly species richness. This effect is similar to the effect size found based on the traditional linear models (Fig. 1). The direct effect of N deposition on butterfly species richness, which is independent from vegetation structure and plant species richness, might be caused by N deposition resulting in reduced food plant quality. While we are not aware of other explanations that could convincingly explain a direct negative effect of N deposition on butterfly species richness, it seems nevertheless unlikely that high N deposition reduces food plant quality so much that this reduces the number of butterfly species considerably.

### Other drivers of butterfly species richness: importance of microclimate

In our systematic search for studies that compared different drivers of butterfly species richness, we found that vegetation variables were much less frequently investigated than environmental or habitat variables. However, if the effects of vegetation variables on butterfly species richness were studied, they were usually described as relevant and consistent. The results of our literature review thus suggest that vegetation variables representing microclimate or plant resource diversity are important but underrepresented in published research on the spatial variation of butterfly species richness.

Our results from structural equation models (SEM) applied to the BDM data confirm the importance of vegetation variables: butterfly species richness was correlated with plant species richness to a similar degree as with ambient temperature. In contrast, the observed effects of land-use intensity and habitat availability were rather weak. An explanation might be that the available information about land-use intensity at the study plots of the BDM was limited. Our predictor variables are mainly derived from the plant surveys and contain average indicator values per 1-km^2^ study plot, therefore hiding within-site variability.

## Conclusions

Whereas in the published literature, N deposition was rarely considered as a driver of butterfly species richness, we found that in Swiss landscapes, high N deposition was consistently linked with low butterfly diversity and low butterfly abundance, suggesting a net loss of butterfly diversity caused by increased N deposition. In addition to agricultural intensity, habitat fragmentation and climate change, atmospheric nitrogen deposition might thus play an essential, yet apparently underestimated, role in threatening butterfly diversity and abundance.

## Supporting information

Web of Science search setting used for the systematic literature review.

Excel file with a table listing the 32 studies used in the literature review and a table listing the predictor variables extracted from these studies.

Matrix of scatterplots between all predictor variables. The given numbers refer to the correlation coefficient of the two respective variables.

Path diagram of the generic model that we used as a starting point for the analysis using structural equation models. Observed variables are depicted

Excel file listing the butterfly species that were recorded in at least 20 study plots. For each species, the estimated effect size of N deposition on

Results of the structural equation model that allows for a direct effect of Nitrogen deposition on butterfly species richness.

## Acknowledgments

We thank the skilled botanists and entomologists who conducted the fieldwork. The Swiss Federal Office for the Environment (FOEN) kindly provided biodiversity monitoring data and topographic data. This work was supported by the FOEN to L.K., B.R. and T.R. and by the Swiss National Science Foundation (grant no. 156294) and the MAVA foundation to V.A.

## Supporting Information

**Figure A1:** Web of Science search setting used for the systematic literature review.

Webofscience_Search_Setting.png

**Table A2:** Excel file with a table listing the 32 studies used in the literature review and a table listing the predictor variables extracted from these studies.

**Figure A3:** Matrix of scatterplots between all predictor variables. The given numbers refer to the correlation coefficient of the two respective variables.

**Figure A4:** Path diagram of the generic model that we used as a starting point for the analysis using structural equation models. Observed variables are depicted in rectangles; latent variables that were measured using several of the predictor variables are depicted in ovals. Arrows indicate assumed causal relationships between variables.

**Table A5:** Excel file listing the butterfly species that were recorded in at least 20 study plots. For each species, the estimated effect size of N deposition on the abundance (number of recorded individuals) is given.

**Figure A6:** Results of the structural equation model that allows for a direct effect of Nitrogen deposition on butterfly species richness. The thickness of the arrows is relative to the absolute values of the effect sizes, with arrows in grey indicating positive effects, and arrows in orange indicating negative effects.

