## Supplementary material for "Nitrogen deposition is negatively related to species richness and abundance of threatened species in Swiss butterflies": Web of Science search setting used for the systematic literature review.

| | | |
|---|---|---|
| | butterfl* OR lepidoptera | Title |
| And | diversity OR richness | Title |
| And | "global change" OR driver* OR predictor* OR variable* | Topic |
| Not | island OR *tropic* | Title |
