## Supplementary figures and images for "Nitrogen deposition is negatively related to species richness and abundance of threatened species in Swiss butterflies"

### Matrix of scatterplots between all predictor variables. The given numbers refer to the correlation coefficient of the two respective variables.

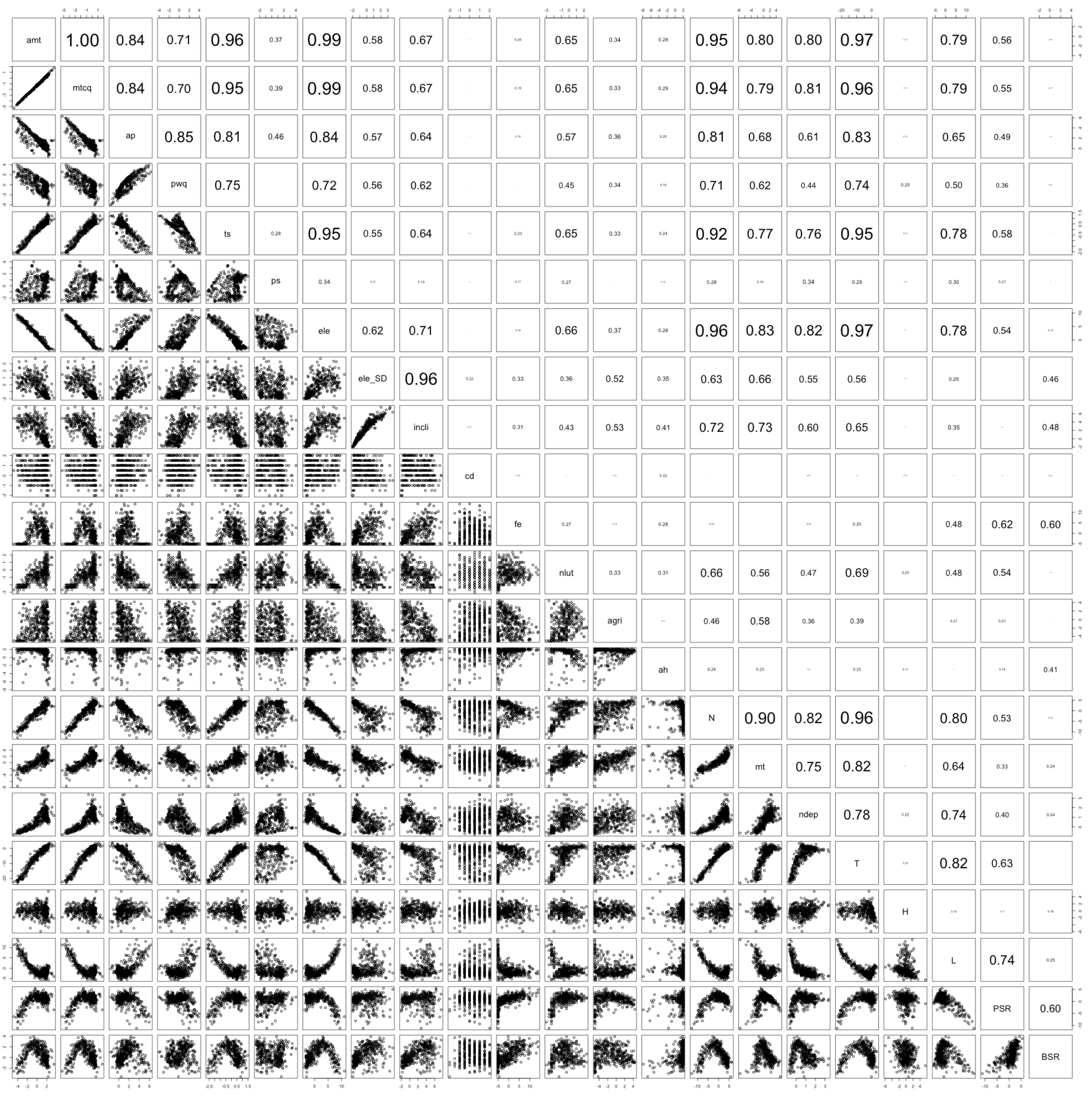

### Path diagram of the generic model that we used as a starting point for the analysis using structural equation models. Observed variables are depicted

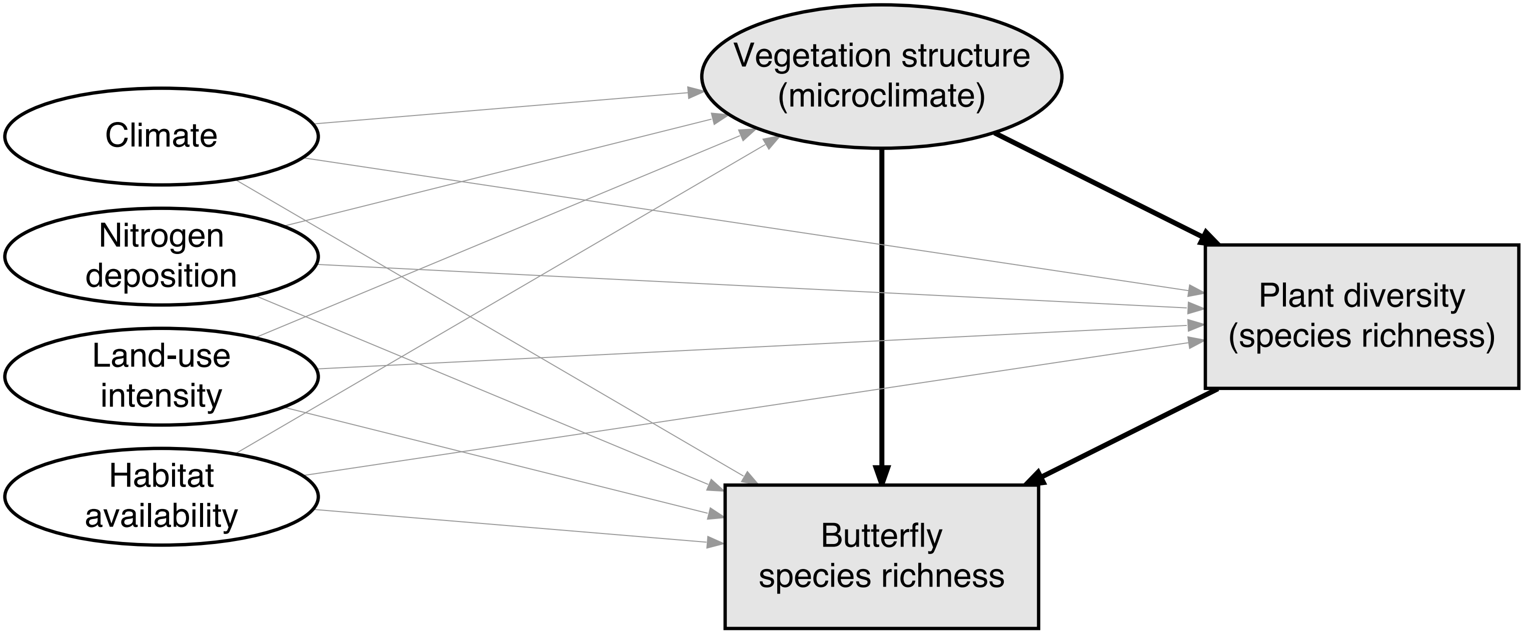

### Results of the structural equation model that allows for a direct effect of Nitrogen deposition on butterfly species richness.

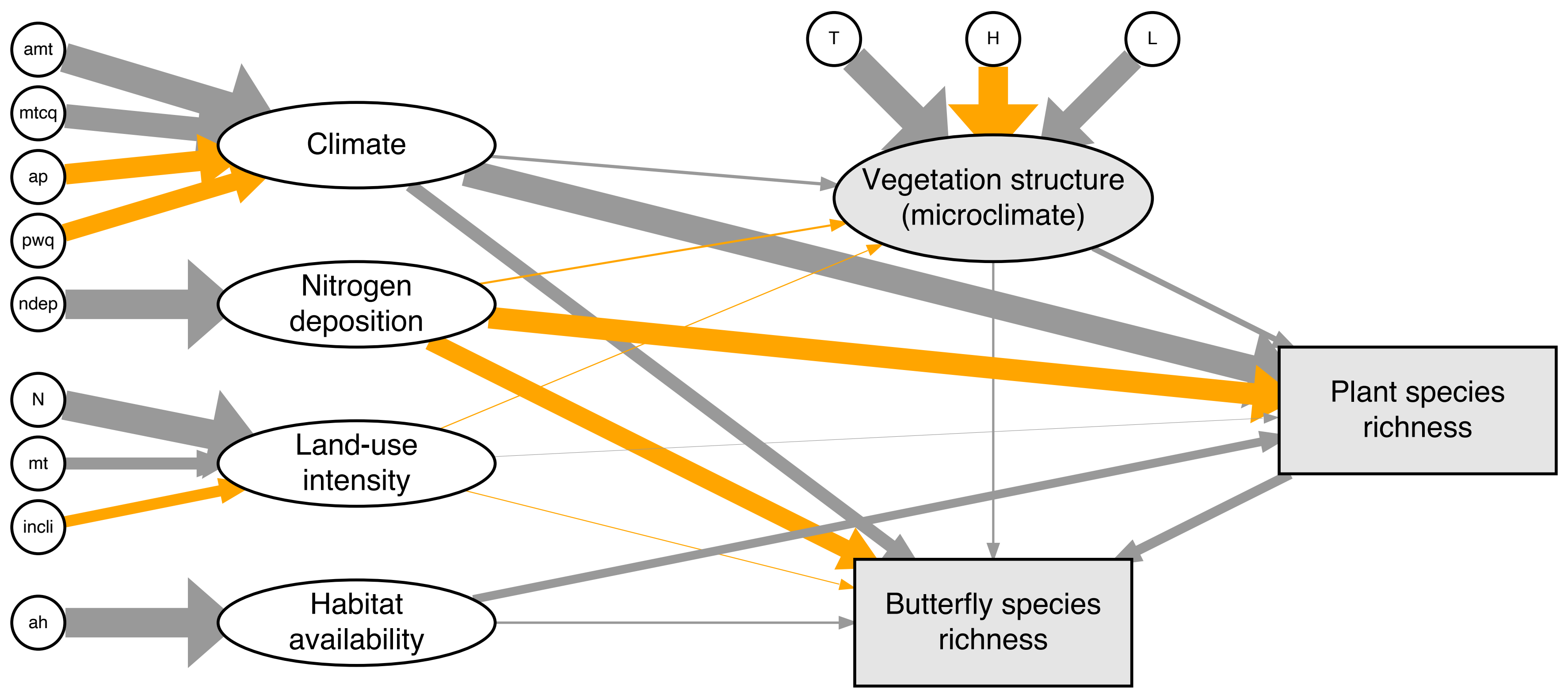
